# Functional and anatomical connectivity predict brain stimulation’s mnemonic effects

**DOI:** 10.1101/2023.07.27.550851

**Authors:** Youssef Ezzyat, James E. Kragel, Ethan A. Solomon, Bradley C. Lega, Joshua P. Aronson, Barbara C. Jobst, Robert E. Gross, Michael R. Sperling, Gregory A. Worrell, Sameer A. Sheth, Paul A. Wanda, Daniel S. Rizzuto, Michael J. Kahana

## Abstract

Closed-loop direct brain stimulation is a promising tool for modulating neural activity and behavior. However, it remains unclear how to optimally target stimulation to modulate brain activity in particular brain networks that underlie particular cognitive functions. Here, we test the hypothesis that stimulation’s behavioral and physiological effects depend on the stimulation target’s anatomical and functional network properties. We delivered closed-loop stimulation as 47 neurosurgical patients studied and recalled word lists. Multivariate classifiers, trained to predict momentary lapses in memory function, triggered stimulation of the lateral temporal cortex (LTC) during the study phase of the task. We found that LTC stimulation specifically improved memory when delivered to targets near white matter pathways. Memory improvement was largest for targets near white matter that also showed high functional connectivity to the brain’s memory network. These targets also reduced low-frequency activity in this network, an established marker of successful memory encoding. These data reveal how anatomical and functional networks mediate stimulation’s behavioral and physiological effects, provide further evidence that closed-loop LTC stimulation can improve episodic memory, and suggest a method for optimizing neuromodulation through improved stimulation targeting.

## Introduction

Direct electrical stimulation of the human brain can manipulate circuits underlying perception, cognition, and action (Siddiqi et al., 2022; Scangos et al., 2021b). Such stimulation has been used to treat network syndromes of brain dysfunction, suggesting that stimulation influences a broader network of brain regions beyond the stimulated location (Mayberg et al., 2005; Scangos et al., 2021a; Limousin et al., 1998; Bouthour et al., 2019; Deuschl et al., 2006; Geller et al., 2017; Jobst et al., 2017; Lozano and Lipsman, 2013). Stimulation can also modulate behaviors, such as learning and memory, that depend on the coordinated activity of a network of brain regions (Mankin and Fried, 2020; Das and Menon, 2021; Voytek and Knight, 2015; Keerativittayayut et al., 2018; Staresina and Wimber, 2019).

Although increasingly used as a therapeutic and experimental tool, variability in outcomes poses a critical challenge, in part because stimulation’s mechanisms of action remain poorly understood. Theoretical accounts evolved from models of local disruption of pathological activity (Benabid et al., 2004) to modulation of the broader network of areas connected to the stimulated location (Ashkan et al., 2017; McIntyre and Hahn, 2010). If stimulation’s effects are best understood at the network level, perhaps variability in individual network structure can explain the variability in physiological and behavioral outcomes.

In support of this idea, applying stimulation to gray matter, the gray-white matter boundary, or specific white matter fibers determines the spread of physiological effects through the network (Paulk et al., 2022; Solomon et al., 2018). Compared to gray matter stimulation, white matter stimulation leads to more broadly distributed excitation in downstream areas (Crocker et al., 2021; Paulk et al., 2022; Mohan et al., 2020). White matter pathways also constrain stimulation’s down-stream functional effects (Lujan et al., 2013; Khambhati et al., 2019; Stiso et al., 2019). Behaviorally, stimulation of white matter has led to remission in depression (Mayberg et al., 2005), slowed cognitive decline in Alzheimer’s (Lozano et al., 2016; Hamani et al., 2008), and enhanced memory in epilepsy (Suthana et al., 2012; Mankin et al., 2021; Titiz et al., 2017).

In addition to the brain’s anatomical architecture, research shows that functional architecture also mediates the spread and persistence of stimulation’s physiological effects (Keller et al., 2011; Fox et al., 2020; Fox et al., 2014; Keller et al., 2018). Previous work further suggests this relation to be frequency-specific. For example, stimulating targets in the medial temporal lobe leads to greater downstream changes in low-frequency (5-13 Hz) activity in brain regions that are strongly connected, at low-frequencies, to the stimulated site (Solomon et al., 2018). There are a variety of cognitive functions, including episodic memory, that have been linked to modulation of low-frequency activity (Colgin, 2013; Burke et al., 2013; Solomon et al., 2017; Donoghue et al., 2020; Koster and Gruber, 2022; Griffiths et al., 2021). Therefore, these physiological findings suggest that stimulating targets with strong low-frequency network connectivity could reliably modulate such behaviors, although this idea has yet to be tested.

We hypothesized that anatomical and functional characteristics of the stimulation target represent key variables that control the effect of stimulation on the brain’s memory network. We applied stimulation in closed-loop in 47 patients while they participated in an episodic memory task (free recall). We stimulated 57 targets located in the lateral temporal cortex (LTC), with the timing of stimulation determined by multivariate classification of neural activity during the encoding phase of the memory task. Using patient-specific data, we characterized each stimulation target based on its proximity to the nearest white matter pathway, as well as its low-frequency resting-state functional connectivity with the the brain’s memory-encoding network. We found that closed-loop LTC stimulation improves memory performance relative to random stimulation, extending prior evidence that LTC stimulation modulates episodic memory (Ezzyat et al., 2018; Kucewicz et al., 2018). Further, we reveal that stimulation target proximity to white matter and functional connectivity predict both stimulation’s effects on memory performance and changes in rhythmic low-frequency activity involved in successful memory encoding.

## Experimental Procedures

### Participants

Forty-seven patients undergoing intracranial electroencephalographic monitoring as part of clinical treatment for drug-resistant epilepsy were recruited to participate in this study. In total, *N* = 57 brain locations were stimulated: 38 patients were stimulated in one location, 8 patients were stimulated in two separate locations, and 1 patient was stimulated in three separate locations. Only one location was stimulated per session. Of the current dataset, data from 14 patients were included in an earlier publication (Ezzyat et al., 2018). All of the presently reported analyses and results are novel.

Data were collected as part of a multi-center project designed to assess the effects of electrical stimulation on memory-related brain function. Data were collected at the following centers: University of Texas Southwestern Medical Center (Dallas, TX), Dartmouth-Hitchcock Medical Center (Lebanon, NH), Thomas Jefferson University Hospital (Philadelphia, PA), Emory University Hospital (Atlanta, GA), Mayo Clinic (Rochester, MN), Hospital of the University of Pennsylvania (Philadelphia, PA), and Columbia University Medical Center (New York, NY). The research protocol was approved by the IRB at each hospital and informed consent was obtained from each participant. Electrophysiological data were collected from electrodes implanted subdurally (grid/strip configurations) on the cortical surface and/or electrodes within the brain parenchyma (depth electrodes). The clinical team determined the placement of the electrodes based on the epileptogenic monitoring needs of the patient.

### Anatomical localization

Cortical surface regions were delineated on pre-implant whole brain volumetric T1-weighted MRI scans using Freesurfer (Fischl et al., 2004) according to the Desikan-Kiliany atlas (Desikan et al., 2006). Whole brain and high resolution medial temporal lobe volumetric segmentation was also performed using the T1-weighted scan and a dedicated hippocampal coronal T2-weighted scan with Advanced Normalization Tools (ANTS) (Avants et al., 2008) and Automatic Segmentation of Hippocampal Subfields (ASHS) multi-atlas segmentation methods (Yushkevich et al., 2015). Coordinates of the radiodense electrode contacts were derived from a post-implant CT and then registered with the MRI scans using ANTS. Subdural electrode coordinates were further mapped to the cortical surfaces using an energy minimization algorithm (Dykstra et al., 2012). Two neuro-radiologists reviewed cross-sectional images and surface renderings to confirm the output of the automated localization pipeline. Stimulation targets that were localized to the left inferior, middle, and superior temporal gyri were classified as LTC. For region of interest analyses, electrodes were assigned to regions using Freesurfer atlas labels (IFG: inferior frontal gyrus; MFG: middle frontal gyrus; SFG: superior frontal gyrus; MTLC: medial temporal lobe cortex; HIPP: hippocampus; ITG: inferior temporal gyrus; MTG: middle temporal gyrus; STG: superior temporal gyrus; IPC: inferior parietal cortex; SPC: superior parietal cortex; OC: occipital lobe).

### Verbal memory task

Across participants, data were collected from two behavioral tasks: standard delayed free recall and categorized delayed free recall. In both tasks, participants were instructed to study lists of words for a later memory test; no explicit encoding task was used. Lists were composed of 12 words presented in either English or Spanish, depending on the participant’s native language. In the standard free recall task, words were selected randomly from a pool of common nouns (https://memory.psych.upenn.edu/Word_Pools). In the categorized free recall task, the word pool was constructed from 25 semantic categories (e.g. fruit, furniture, office supplies). Each list of 12 items in the categorized version of the task consisted of four words drawn from each of three categories. Overall, *N* = 19 participated in standard free recall only; *N* = 26 participated in categorized free recall only; and *N* = 2 participated in both free and categorized recall (in separate sessions).

Immediately following the final word in each list, participants performed a distractor task (to attenuate the recency effect in memory, length = 20 seconds) consisting of a series of arithmetic problems of the form A+B+C=??, where A, B and C were randomly chosen integers ranging from 1-9. Following the distractor task participants were given 30 seconds to verbally recall as many words as possible from the list in any order; vocal responses were digitally recorded and later manually scored for analysis. Each session consisted of 25 lists of this encoding-distractor-recall procedure.

### EEG recording and analysis

Electrophysiological recording and stimulation was conducted using a variety of systems across the sites over which the project was conducted. Recording and stimulation equipment included clinical EEG systems (Nihon Kohden EEG-1200, Natus XLTek EMU 128 or Grass Aura-LTM64), equipment from Blackrock Microsystems, as well as the External Neural Stimulator (ENS) (Medtronic, Inc.). Data were sampled at 500, 1000, or 1600 Hz (depending on the clinical site). During the sessions, a laptop recorded behavioral responses (vocalizations, key presses), synchronized to the recorded EEG via transmitted network packets.

Intracranial electrophysiological data were filtered to attenuate line noise (5 Hz band-stop fourth order Butterworth, centered on 60 Hz). We referenced the data using a bipolar montage (Burke et al., 2013) by identifying all pairs of immediately adjacent contacts on every depth, strip and grid and taking the difference between the signals recorded in each pair. The resulting bipolar timeseries was treated as a virtual electrode and used in all subsequent analysis. For the purposes of anatomical localization, we used the midpoint of the bipolar pair as the location for this virtual electrode. We used the same midpoint approach to localize stimulation targets and to measure stimulation target distance to white matter (see below).

### Multivariate classification of memory

We performed spectral decomposition (8 frequencies from 6-175 Hz, logarithmically-spaced; Morlet wavelets; wave number = 5) for 1366 ms epochs from 0 to to 1366 ms relative to word onset. Mirrored buffers (length = 1365 ms) were included before and after the interval of interest to avoid convolution edge effects. The resulting time-frequency data were then log-transformed, averaged over time, and *z*-scored within session and frequency band across word presentation events. For a subset of participants, we also performed the same spectral decomposition procedure on record-only data from the memory recall phase of each list. These data were then used in addition to the encoding data to train the classifier (Kragel et al., 2017). To do so, we computed spectral power for the 500 ms interval preceding a response vocalization, as well as during unsuccessful periods of memory search (the first 500 ms of any 2500 ms interval in which no recall response was made). For both trial types (correct vocalizations and unsuccessful search periods), we further stipulated that no vocalization onsets occurred in the preceding 2000 ms.

Our closed-loop stimulation approach was based on using individualized memory classifiers to control the timing of stimulation in response to brain activity. Thus, after collecting at least three record-only sessions from an individual patient, we then used the data as input to a logistic regression classifier that would trigger closed-loop stimulation during the later stimulation session(s). To build the classifier, we used patterns of brain activity collected during record-only sessions and trained the classifier to discriminate words that were recalled vs. not recalled. The input features were spectral power at the eight analyzed frequencies × *N* electrodes (Fig 1A). We used L2-penalization to prevent overfitting (Hastie et al., 2001) and set the penalty parameter (*C*) to 2.4 × 10^−4^ based on the optimal penalty parameter calculated across our large pre-existing dataset of free-recall participants (Kragel et al., 2017; Ezzyat et al., 2018). We weighted the penalty parameter separately for each participant in inverse proportion to their number of recalled and not recalled words; this was done so that the model would learn equally from both classes (Hastie et al., 2001). Classification analyses were programmed using either the Matlab implementation of the LIBLINEAR library (Fan et al., 2008) or the Python library scikit-learn (Pedregosa et al., 2011). For the Closed-loop group (34 participants, *N* = 40 stimulation targets), classifiers were trained using the true mapping of features (spectral power × electrodes) to recall outcomes. In contrast, for the Random group (13 participants, *N* = 17 stimulation targets), a technical error in labeling features during classifier training led to classifiers that were trained on permuted data, eliminating the true mapping between neural activity on each trial and recall outcomes. This provided a natural experiment for testing whether the closed-loop nature of stimulation enhanced the efficacy of LTC stimulation.

**Figure 1:**
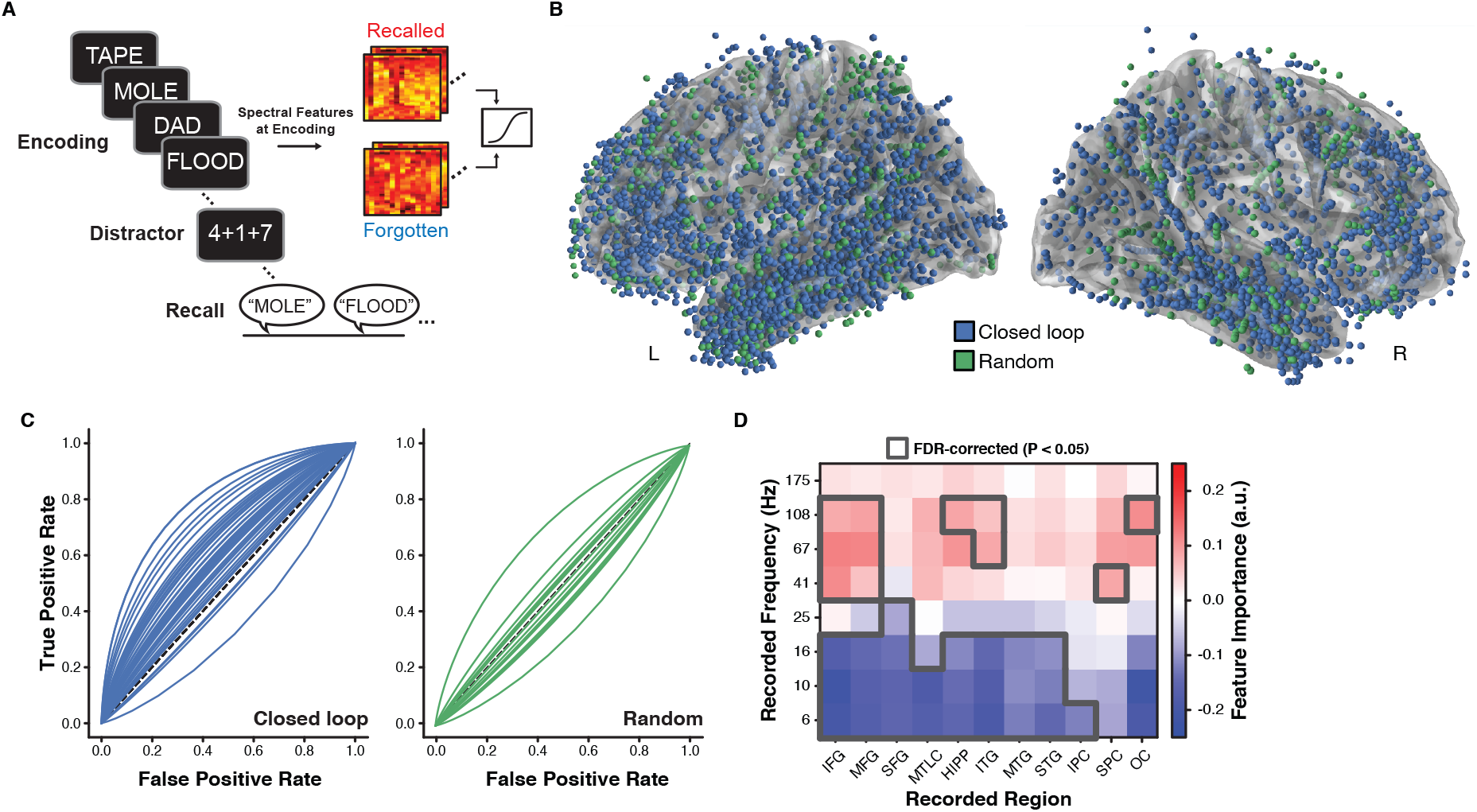
Stimulation strategy and classifier performance. **A.** Participants performed at least three sessions of the free recall task while being monitored with intracranial EEG. Multivariate classifiers trained on whole-brain patterns of spectral activity predicted subsequent recalled vs. not recalled words. **B.** Recording electrode locations for all participants in the Closed-loop (blue) and Random (green) groups, rendered on the Freesurfer average brain. **C.** Each participant’s multivariate classifier then served as their personalized model to trigger stimulation. Classifiers trained on record-only data generalized to the stimulation session(s) for the Closed-loop group (*P* = 6.14 × 10^−7^) and outperformed classifiers for the Random group (*P* = 2.73 × 10^−5^). **D.** An analysis of feature importance for classifiers from the Closed-loop group showed that successful memory states were associated with decreases in low-frequency activity and increases in high-frequency activity.

To assess the importance of individual features to the classifier’s performance, we calculated a forward model (Haufe et al., 2014):

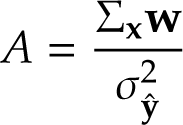

where Σ**_x_** is the data covariance matrix, **w** is the vector of feature weights from the trained classifier, and σ^2^ is the variance of the logit-transformed classifier outputs for all recalled/not recalled events **y**^. Positive values in *A* suggest a positive relation between power for a given feature and successful memory recall. We computed *A* separately for each participant (averaging features within anatomical regions of interest based on the Freesurfer labels derived from anatomical localization of electrodes) before conducting across-participant statistical tests (Fig 1D).

### Closed-loop stimulation

At the start of each stimulation session, we determined the safe amplitude for stimulation using a mapping procedure in which stimulation was applied at 0.5 mA while a neurologist monitored for afterdischarges. This procedure was repeated, incrementing the amplitude in steps of 0.5 mA, up to a maximum of 1.5 mA for depth contacts and 3.5 mA for cortical surface contacts. These maximum amplitudes were chosen to be below the afterdischarge threshold and below accepted safety limits for charge density (Shannon, 1992). For each stimulation session, we passed electrical current through a single pair of adjacent electrode contacts. The locations of implanted electrodes were determined strictly by the monitoring needs of the clinicians (recording sites depicted in Figure 1B). We therefore used a combination of anatomical and functional information to select stimulation sites, prioritizing (if available) targets in the middle temporal gyrus (stimulation targets depicted in Figure 2A). This choice was guided by prior work identifying the middle temporal gyrus as an effective target for modulating memory with stimulation (Ezzyat et al., 2018; Kucewicz et al., 2018). Stimulation was delivered using charge-balanced biphasic rectangular pulses (pulse width = 300 µs) at either 50, 100 or 200 Hz frequency (a single frequency was chosen for each subject), and was applied for 500 ms in response to classifier-detected poor memory states (see below). Participants performed one practice list followed by 25 task lists: lists 1-3 were used as a baseline for normalizing the classifier; lists 4-25 consisted of 11 lists each of Stim and NoStim conditions, randomly interleaved. NoStim lists were identically structured to Stim lists, except that stimulation was never delivered in response to classifier output.

**Figure 2:**
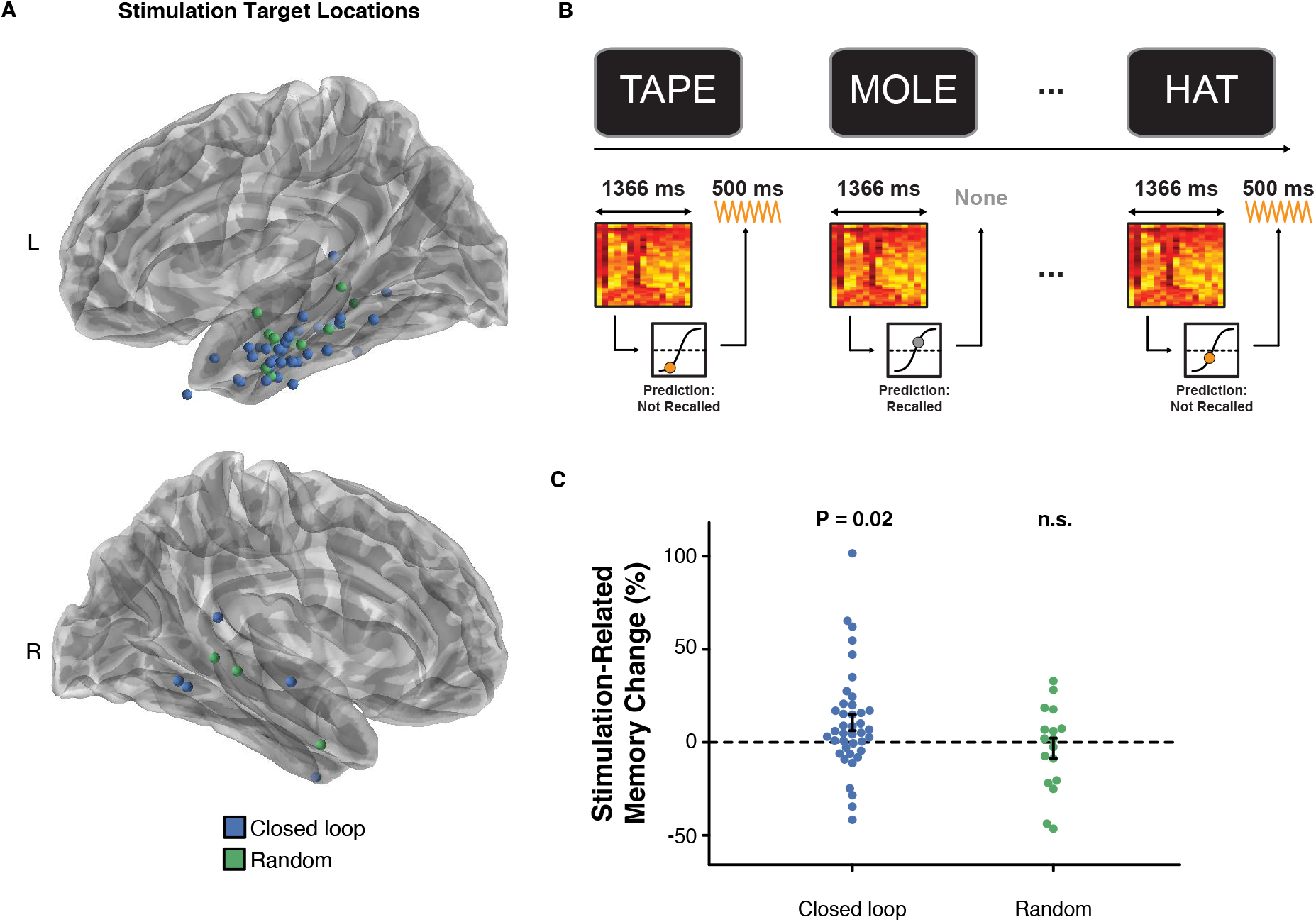
Closed-loop stimulation improves memory performance. **A.** Stimulation target locations for the Closed loop (blue) and Random (green) groups. **B.** Closed-loop stimulation strategy. **C.** Closed-loop LTC stimulation improved memory performance (*P* = 0.02) while Random stimulation did not. Error bars in *C* reflect s.e.m. Closed-loop and Random groups [Mann-Whitney *U* = 331.0, *P* = 0.44] and the median distance was in fact numerically greater for the Closed loop (1.58 mm) compared to the Random group (1.39 mm). This suggests that distance to white matter alone does not explain the finding of improved memory in the Closed-loop group. Instead, proximity to white matter appears to enhance the effectiveness of closed-loop stimulation.

To determine (in actuality) how well the classifier predicted recalled and forgotten words in a given participant’s stimulation session, we again used AUC. We used the true classifier outputs and true recall outcomes from the NoStim lists to calculate the classifier generalization AUC for the stimulation sessions. To generate the corresponding receiver operating characteristic curves for visualization (Figure 1C), we modeled the classifier outputs for recalled and not recalled words using signal detection theory (Wixted, 2007). We did this by using the classifier outputs to estimate the mean and variance of hypothetical (normal) distributions of memory strength for recalled and not recalled words. We then generated a curve relating true and false positive rates by varying the assumed decision criterion (Wixted, 2007).

### Analysis of memory performance

All participants completed at least three sessions of the record-only task (for purposes of classifier training) and at least one session of the stimulation task. For the stimulation session(s) we calculated stimulation’s effect on recall performance as follows:

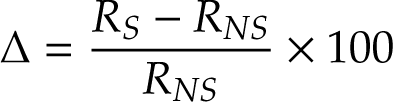

where *R_S_*is the average recall for stimulated lists and *R_NS_* is the average recall for non-stimulated lists. Because the first three lists of every stimulation session were always non-stimulated (used for normalization of the classifier input features for that session), we excluded these lists from the calculation of *R_NS_* to avoid introducing a temporal order confound (Ezzyat et al., 2018). All participants were required to demonstrate a minimum *R_NS_* = 8.33% (1 out of 12 words per list) for inclusion in the sample.

### Calculation of stimulation target distance to white matter

Using Freesurfer to segment patients’ T1 MRI scan, we identified white-matter vertex locations, then calculated the distance between the stimulation location (midpoint of the bipolar pair) and the nearest white matter vertex. These distances were then split into thirds in order to categorize stimulation sites as Near, Mid, or Far relative to the nearest white matter (Solomon et al., 2018; Mohan et al., 2020).

### Calculation of stimulation target node strength

We adapted a previously reported method for calculating the resting-state functional connectivity between channels using the MNE-Python software package (Gramfort et al., 2014; Solomon et al., 2018). We extracted data from non-task periods of the record-only sessions of each patient and used the data to calculate the coherence between each pair of bipolar channels in the patient’s montage. The coherence (*C_xy_*) between two signals is the normalized cross-spectral density. This measure reflects the consistency of phase differences between signals at two electrodes, weighted by the correlated change in spectral power at both sites:

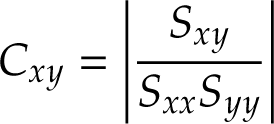

where *S_xy_* is the cross-spectral density between signals at electrodes *x* and *y*; *S_xx_* and *S_yy_*are the auto-spectral densities at each electrode. We used the multitaper method to estimate spectral density (Bronez, 1992). We used a time-bandwidth product of 4 and a maximum of 8 tapers (tapers with spectral energy < 0.9 were removed), computing coherence for frequencies between 5 and 13 Hz. We computed inter-electrode coherence within non-overlapping 1-s windows of data collected during a 10-second baseline (countdown) period that occurred at the start of each word list. The resulting coherence values between each pair of electrodes were then regressed on the Euclidean distance between each pair of electrodes, to account for the correlation between inter-electrode coherence and distance (Solomon et al., 2018). This distance-residualized measure of coherence was then used in the node-strength calculation. We repeated this entire procedure for calculating high-frequency functional connectivity in the 45-90 Hz range.

### Analysis of physiological e**ff**ects of stimulation

To assess the effect of LTC stimulation on neural activity, we analyzed recording channels (i.e. those that were not stimulated) and we compared stimulation-evoked spectral power separately at low and high frequencies. We first excluded electrodes exhibiting non-physiological post-stimulation artifacts (such as amplifier saturation/relaxation) using three different measures of the EEG timeseries before and after stimulation. We compared intervals before and after stimulation for changes in variance using an *F*-test and for changes in signal amplitude using a *t*-test. We additionally fit a polynomial function to the timeseries before and after each stimulation event and used a *t*-test to compare the resulting parameter estimates for the quadratic term. We calculated these three measures using the signal from -400 ms to -100 ms relative to stimulation onset and 100 ms to 400 ms relative to stimulation offset. In order to select statistical thresholds for each measure, we conducted the same analysis on each participant’s record-only data. We then selected *P*-value thresholds associated with a 5% detection rate in the record-only data (i.e. false positives). Any channel that was significant on any of the three measures was excluded from analysis.

To measure stimulation’s effect on low-frequency power, we extracted spectral power from -600 ms to -100 ms relative to stimulation onset and 100 ms to 600 ms relative to stimulation offset. We used Morlet wavelets (wave number = 5) to estimate spectral power for the same set of frequencies used to train the classifier with buffers to eliminate edge artifacts. The resulting spectral power estimates were then *z*-scored within each frequency, separately for each session. We then averaged power within each frequency across the time dimension for each pre-stimulation period and for each matched post-stimulation period. We then subtracted the pre-stimulation data from the post-stimulation data to yield a distribution of change in spectral power for each electrode.

We compared the distribution of power changes for stimulation events to the power changes from matched intervals on NoStim lists. To do so, we calculated spectral power using identical parameters. However, because there were no actual stimulation events in NoStim lists, we generated a synthetic distribution of stimulation onset times by extracting the lag (in milliseconds) between each word onset and stimulation event in Stim lists, and sampling randomly from that distribution of onset times to determine when to extract data relative to word onset events in NoStim lists.

Finally, we used an independent samples *t*-test to compare the distribution of Stim list power differences to the distribution of NoStim list power differences within each electrode. The resulting distribution of *t*-statistics was then averaged across electrodes to estimate the stimulation-evoked change in power (Figure 5A). We then averaged these values separately within clusters of low and high frequencies that significantly predicted memory performance (based on classifier feature importance, Figure 1D).

### Statistics

Data are presented as mean ± standard error of the mean; scatterplots show the standard error of the estimate. All statistical comparisons were conducted as two-tailed tests. Non-parametric tests (Mann-Whitney; Spearman rank correlation) were used for non-normally distributed variables (e.g. white matter distance, Figure 3A); parametric tests (*t*-tests and Pearson correlation) were used for the remaining analyses. Data distributions were visually inspected or assumed to be normal for parametric tests.

**Figure 3:**
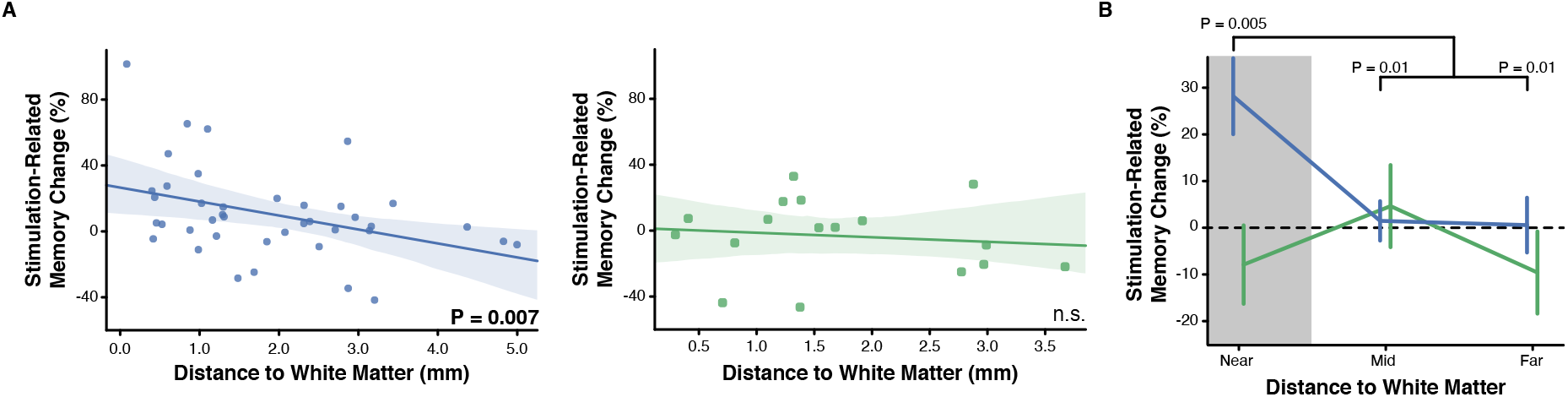
Closed-loop stimulation near white matter enhances memory. **A.** For the Closed-loop group, stimulation’s effect on memory depended on the target distance from the nearest white matter [*left*, *P* = 0.007]. The correlation was not significant for the Random group [*right*, *P* = 0.52]. **B.** Closed-loop LTC stimulation improved memory performance for targets located nearest to white matter (*P* = 0.005). There was no effect for the Random group (*P* = 0.62). Error regions in *A* reflect the standard error of the estimate. Error bars in *B* reflect s.e.m.

### Data Availability

Upon publication, all de-identified raw data and analysis code may be downloaded at http://memory.psych.upenn.edu/Electrophysiological_Data.

## Results

### Multivariate classifiers identify memory lapses

Our stimulation strategy sought to intercept and rescue periods of poor memory encoding. To do so, we trained participant-specific multivariate classifiers to discriminate patterns of neural activity during record-only sessions of free recall (Figure 1A). For the Closed-loop group (*N* = 40), classifiers were trained using the true mapping of features (spectral power × electrodes) to recall performance; for the Random group (*N* = 17), due to a technical error in labeling features (see *Methods*), classifiers were trained on permuted features. The recording electrode locations for the Closed-loop and Random groups appear as spheres in Figure 1B. After training the classifiers on record-only data, we used them in later (independent) sessions to identify poor memory states for targeting with stimulation.

Our first question was how well the classifiers predicted memory outcomes during the stimulation sessions (i.e. out-of-sample generalization). To answer this question, we used data from NoStim lists in which we obtained classifier predictions about the probability of recall for each word, but did not use these predictions to trigger stimulation (see *Methods*). Using area under the receiver operating characteristic curve as an index of classification accuracy, we found that classifiers for the Closed-loop reliably exceeded chance performance [Mean AUC = 0.62 (chance AUC = 0.50), Wilcoxon signed rank test *P* = 5.73 × 10^−7^]. Closed-loop classifiers also outperformed classifiers for the Random group [Mann-Whitney *U* = 586.0, *P* = 1.85 × 10^−5^]. As expected, Random classifiers did not exceed chance [Mean AUC = 0.49, Wilcoxon signed rank test *P* = 0.55; Figure 1C].

To understand what features the classifier used to discriminate good vs. poor memory encoding states, we used a forward model for each participant to derive importance estimates for each feature (Haufe et al., 2014). We averaged the feature importance values within a set of regions of interest (ROIs) separately for each classifier frequency. Across participants, classifiers predicted successful memory encoding based on increased high-frequency activity (especially in frontal, lateral temporal, and medial temporal lobe areas) and decreased low-frequency activity across much of the recorded cortex and subcortex (Figure 1D). This pattern, which we refer to as the spectral tilt, has been observed in previous studies to be a biomarker of successful episodic memory encoding and retrieval (Ezzyat et al., 2017; Burke et al., 2014; Long et al., 2014).

### Closed-loop LTC stimulation improves memory

Having established that classifiers in the Closed-loop group reliably discriminate memory encoding states, we next asked if we could increase memory performance via stimulation of the LTC (Figure 2A). Our stimulation strategy was based on detecting poor memory encoding states and intercepting them with stimulation (Figure 2B). For the Closed-loop and Random groups, we compared recall performance for lists in which we delivered stimulation (Stim lists) vs. identically structured lists in which we did not stimulate (NoStim lists, as described above). In the Closed-loop group, recall was higher on Stim lists compared to NoStim lists [Δ = 10.6% ± 4.3; *t*(39) = 2.45, *P* = 0.02, Figure 2C], suggesting that intercepting poor memory encoding states with LTC stimulation enhanced recall. In contrast, there was no difference in memory performance for the Random group [Δ = −3.2% ± 5.5; *t*(16) = −0.59, *P* = 0.57]. There was a trend for greater memory enhancement for the Closed-loop compared to the Random group [*t*(55) = 1.83, *P* = 0.07]. These findings are the first to directly compare closed-loop LTC stimulation with a random/open-loop stimulation control and, in a larger replication sample, demonstrate the robustness of previous studies showing memory enhancement via LTC stimulation (Ezzyat et al., 2018; Kucewicz et al., 2018; Kahana et al., 2023).

### White matter proximity mediates stimulation’s e**ff**ect on memory

Motivated by physiological studies of electrical stimulation’s effects on downstream targets (Mohan et al., 2020; Solomon et al., 2018; Keller et al., 2018), we asked whether stimulating close to white matter tracts would produce greater positive or negative effects on memory. If so, this would suggest that the brain’s anatomical network structure plays a key role in determining how effectively stimulation can modulate cognitive function (Stiso et al., 2019; Crocker et al., 2021). To answer this question, we examined how stimulation’s effect on memory performance varied as a function of the stimulation target’s proximity to white matter (see Figure 3A). For the Closed-loop group, lower distance to white matter predicted greater stimulation-related memory improvement [Spearman ρ(38) = −0.42, *P* = 0.007; Figure 2B]. In the random stimulation group, we neither expected nor observed a correlation between white matter distance and the memory effect [Spearman ρ(15) = −0.17, *P* = 0.52]. There was no difference between the distances to white matter for the To further test this idea, we divided stimulation targets into terciles and asked whether stimulation near white matter was particularly effective in modulating memory performance. Indeed, Closed-loop stimulation targets near white matter enhanced memory performance on Stim lists compared to NoStim lists [Near: *M* = 28.25% ± 8.14%, *t*(13) = 3.23, *P* = 0.005]. This memory improvement was larger than for Closed-loop stimulation targets further away from white matter [Mid: *M* = 1.55% ±4.22%, *P* = 0.01; Far: *M* = 0.64% ±5.87%*P* = 0.01]. Closed-loop stimulation near white matter also significantly outperformed Random stimulation near white matter group [Random *M* = −7.89% ± 8.40%, *t*(17) = 2.36, *P* = 0.03, Figure 2C]. As expected, the Random group did not show improved memory (Stim vs. NoStim within-participant) in any white matter distance bin (all *P* > 0.38). These data suggest that stimulating near white matter leads to greater modulation of memory, and extend previous work that linked white matter proximity to stimulation’s effect on electrophysiology (Mohan et al., 2020; Solomon et al., 2018; Keller et al., 2018; Stiso et al., 2019; Crocker et al., 2021).

### Stimulation target functional connectivity predicts the change in memory

We next asked why closed-loop stimulation delivered near white matter reliably modulated memory function. One possibility is that stimulating near white matter allows more reliable and direct access to the broader memory network connected to the stimulated location (Khambhati et al., 2019; Stiso et al., 2019; Solomon et al., 2018; Mohan et al., 2020). We therefore measured functional connectivity between the brain’s memory encoding network and the stimulation targets located near white matter. Critically, we constructed separate measurements of connectivity at low (5-13 Hz) and high frequencies (45-90 Hz) by calculating coherence using participant-specific resting-state data (see *Methods*). Then, to isolate the brain’s memory encoding network, we identified all electrodes that were in brain regions that showed a spectral tilt that predicted memory success during the task, assessed using classifier feature importance Figure 3A. We then compared stimulation target connectivity to electrodes In vs. Out of the memory network, for both low and high-frequency coherence (referred to as Node Strength). Stimulation targets showed stronger low-frequency connectivity to electrodes in the memory network than to electrodes outside of the memory network [*t*(13) = 3.14, *P* = 0.008, Figure 3B]. For memory network electrodes, low-frequency connectivity was also higher than high-frequency connectivity [*t*(13) = 2.48, *P* = 0.03]. In contrast, stimulation targets showed equivalent high-frequency connectivity In vs. Out of the memory network [interaction: *F*(1, 13) = 10.54, *P* = 0.006, Figure 3B].

Although stimulation targets near white matter showed greater overall low-frequency connectivity with memory-predicting brain areas, this finding leaves open the question of whether variability in connectivity strength with the memory network predicts variability stimulation’s effect on memory. To answer this question, we correlated low-frequency node strength with stimulation-related memory change. We found that low-frequency node strength predicted closed-loop stimulation’s effect on memory [*r*(12) = 0.648, *P* = 0.01, Figure 3C] while high-frequency node strength did not (*P* = 0.65). The difference in correlation for low vs. high-frequency node strength was also significant (two-tailed permutation test *P* = 0.03). For all other targets that were further from white matter, there was no relation between node strength and stimulation-related memory change (all *P* > 0.21).

### Functional connectivity mediates stimulation’s e**ff**ect on downstream physiology

The preceding results indicate that low-frequency functional connectivity to the memory network predicts stimulation effects on memory. Our final question was whether low-frequency connectivity also predicts stimulation’s physiological effects across the memory network. To test this prediction we again examined Closed-loop stimulation targets near white matter and correlated each stimulation target’s connectivity to the memory network with the stimulation-evoked spectral power in this network (Figure 5A). Two participants’ data were excluded due to excessive stimulation artifact on the recording channels. In the remaining participants, we found that stimulation-target functional connectivity predicted stimulation-related changes in low-frequency power [*r*(10) = −0.65, *P* = 0.02, Figure 5B). The correlation was not significant when using high-frequency connectivity and evoked power (*P* = 0.81, Figure 5C).

## Discussion

Direct electrical stimulation has emerged as a powerful tool for manipulating neural activity. The present study evaluated the hypothesis that network properties of a stimulated brain location predict stimulation’s effects on both memory and network physiology. Prior studies suggest that white matter pathways mediate stimulation’s network-level physiological effects (Paulk et al., 2022; Solomon et al., 2018; Mohan et al., 2020; Khambhati et al., 2019; Stiso et al., 2019). Other studies demonstrate that measures of structural and functional connectivity predict stimulation’s effects on downstream targets (Keller et al., 2011; Fox et al., 2020; Solomon et al., 2018). However, none have simultaneously linked structural/functional connectivity with both (1) a reliable improvement over baseline cognitive functioning and (2) concomitant changes in neurophysiology that explain the behavioral effect. To directly address these questions, we asked whether white-matter proximity and functional connectivity underlie the degree to which stimulation of LTC produces improvements or impairments of memory, alongside changes in oscillatory signatures of mnemonic function.

We found that closed-loop stimulation of LTC reliably improved memory on stimulated vs. non-stimulated lists. Consistent with the hypothesis that white-matter pathways convey the effects of stimulation to the broader memory network, we found the benefits of closed-loop LTC stimulation to arise principally from stimulating in, or near, white matter pathways. For the electrodes nearest to white matter, stimulation yielded a 28% increase in recall performance, whereas we failed to observe any reliable increase when delivering stimulation far from these pathways (1%). In a subgroup of subjects who received randomly timed stimulation in LTC targets we failed to observe any improvement in memory performance.

To evaluate how stimulation-–target functional connectivity mediates stimulation’s behavioral and physiological effects, we analyzed participant-specific large-scale neural recordings obtained during prior record-only sessions. Prior studies have shown that brain networks become coherent at low-frequencies during successful memory encoding and retrieval (Solomon et al., 2017; Kragel et al., 2021a), so we used low-frequency coherence to measure the network node strength of each stimulation target. We then asked if greater node strength between LTC stimulation targets and downstream memory-predicting areas resulted in greater effects of stimulation on memory performance. Consistent with this hypothesis, we found a strong positive correlation (*r* = 0.648, see Figure 4C) between low-frequency connectivity and stimulation-related memory improvement. Finally, LTC stimulation engaged low-frequency activity across a broader brain network in a way that matched the network position of the stimulated location (Figure 5).

**Figure 4:**
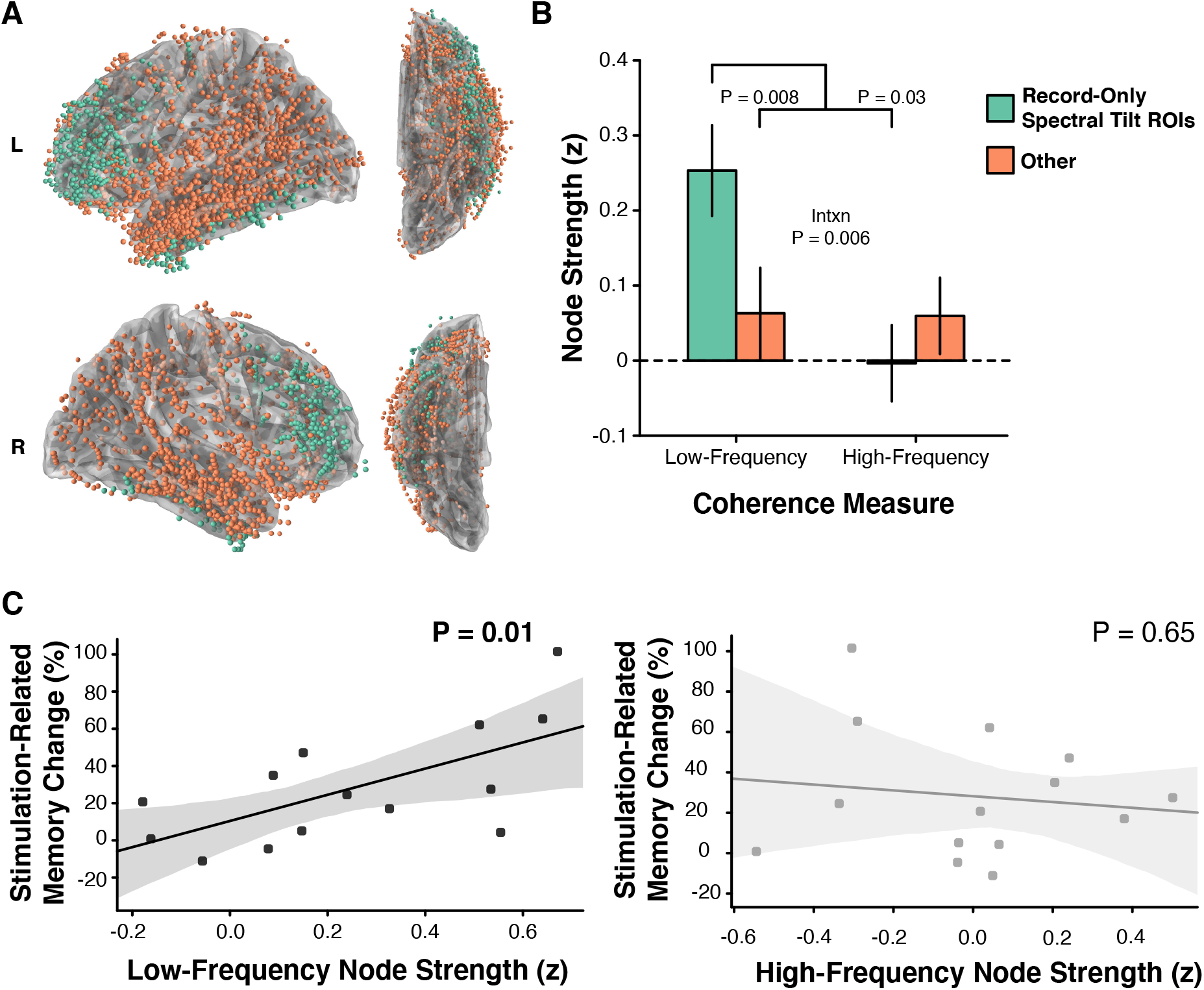
Stimulation target functional connectivity. **A.** We assigned each patient’s record-only electrodes to two ROIs based on whether the electrode was located in a region that showed a memory-related spectral tilt or not (Other). **B.** Low-frequency connectivity was higher between the stimulation target and electrodes in classifier-defined memory regions, compared to electrodes in Other regions (*P* = 0.008) and compared to high-frequency network connectivity (*P* = 0.03). In contrast, there was no difference in stimulation target high-frequency network connectivity. **C.** For closed-loop targets nearest to white matter, there was a significant correlation between stimulation target low-frequency connectivity and stimulation’s effect on memory [*r*(12) = 0.648, *P* = 0.01]. There was no effect for high-frequency connectivity. Errorbars reflect s.e.m. Error regions reflect the standard error of the estimate.

**Figure 5:**
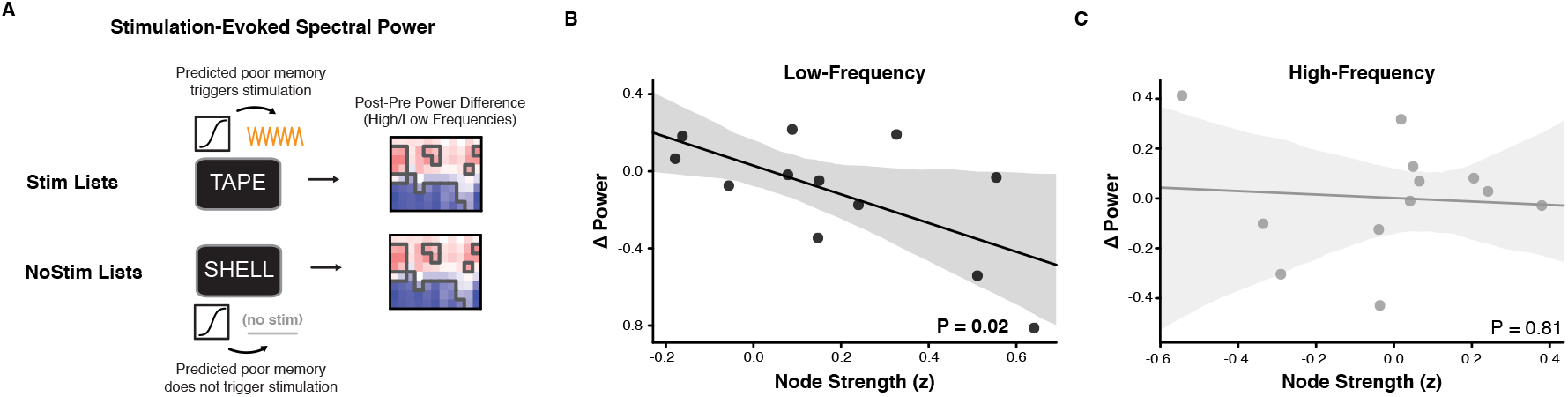
Memory network connectivity predicts physiology. **A.** Schematic of analysis of stimulation-evoked physiology. **B.** For stimulation targets near white matter, low-frequency functional connectivity predicted the stimulation evoked change in low-frequency power (*P* = 0.02). **C.** High-frequency network connectivity did not predict stimulation’s effect on high-frequency activity.

Our data highlight how precise targeting improves stimulation efficacy by showing that delivering stimulation near LTC white-matter leads to greater stimulation-related memory gains (Figure 3C). By linking low-frequency network connectivity with physiological and behavioral outcomes, our study also points to a neural mechanism for modulating memory with stimulation. This result extends earlier work that demonstrated the potential to modulate episodic memory by targeting LTC with stimulation (Ezzyat et al., 2018; Kucewicz et al., 2018). Directly comparing closed-loop and open-loop stimulation strategies in the same study helps to establish a causal role for the closed-loop approach (Hampson et al., 2018; Ezzyat and Rizzuto, 2018). Finally, our data from 57 stimulation targets (across 47 patients) also represents a substantial increase compared to sample sizes described in related prior studies (Ezzyat et al., 2018; Hampson et al., 2018).

Prior work has linked successful memory function with theta power and coherence (Burke et al., 2013; Solomon et al., 2017; Herweg et al., 2020; Griffiths et al., 2019; Kragel et al., 2021b; Ter Wal et al., 2021; Osipova et al., 2006; Guderian and Dü zel, 2005; Klimesch et al., 1997; Staudigl and Hanslmayr, 2013). Here, we investigated this physiological correlate of memory function by testing how memory-modulating LTC stimulation affects low-frequency physiology. We found that stimulation’s effect on low-frequency activity depends on the low-frequency functional connectivity of the stimulation target. This suggests that identifying strong functional connections can produce stronger modulation of low-frequency activity within the memory network. Furthermore, we found that stimulation that modulated low-frequency activity also modulated memory performance.

Several prior studies found potential therapeutic benefits of closed-loop stimulation triggered by decoding of intracranial brain recordings (Ezzyat et al., 2018; Scangos et al., 2021a; Hampson et al., 2018; Kahana et al., 2023). However, with some important exceptions (Hampson et al., 2018), this work has lacked an open-loop or random stimulation control condition, leaving open the question of what *specific* role the closed-loop nature of stimulation played in its therapeutic effects. Here, we compared the effects of closed-loop stimulation with a random stimulation condition. Closed-loop participants received stimulation only for those items predicted to be forgotten. Participants in the random group followed the same protocol, but using classifiers trained on permuted data, resulting in stimulation being applied without regard to predicted memory success. This led to reliable memory improvement for the closed-loop group and none for the random group, despite following an otherwise identical protocol (Figure 1C).

We found that closed-loop stimulation improved memory the most when it was delivered to LTC targets in or near white matter. This finding builds on a growing literature that indicates that stimulation is most effective when it is delivered in or near white matter pathways (Khambhati et al., 2019; Stiso et al., 2019; Mohan et al., 2020; Solomon et al., 2018; Paulk et al., 2022). One explanation for this phenomenon is that only stimulation of white matter pathways successfully engages broader brain networks, perhaps via oscillatory synchronization. In contrast, gray matter stimulation tends to cause more local effects (Mohan et al., 2020; Paulk et al., 2022). Though purely local effects may sometimes be desirable, the key cognitive and pathophysiological processes of greatest interest to neuroscientists tend to involve multiple interconnected brain regions.

Among its many applications for modulating cognition and behavior (Siddiqi et al., 2022; Fox et al., 2020; Sreekumar et al., 2017) a number of recent studies have evaluated stimulation’s potential for enhancing episodic memory (Mankin and Fried, 2020; Suthana and Fried, 2014; Curot et al., 2017; Lee et al., 2013; Sankar et al., 2014). While our study investigated numerous stimulation targets within the LTC, future work should compare stimulation of this region to other brain areas within the broader episodic memory network. Recent work suggests that stimulating white matter pathways in the medial temporal lobe, for example, can also improve memory (Titiz et al., 2017; Mankin et al., 2021; Suthana et al., 2012). However, these previous studies used visual and/or spatial memoranda, while the present study focused on encoding and retrieval of verbal material. Thus, future research should compare stimulation to the lateral and medial temporal lobes, to determine whether stimulation target location interacts with the modality of the to-be-remembered information. This could contribute to other work that has used stimulation to study the component processes that contribute to successful episodic memory (El-Kalliny et al., 2019).

We delivered stimulation using macroelectrodes, consistent with its clinical applications (Krauss et al., 2021; Morrell, 2011; Sun et al., 2008). Macroelectrode stimulation alters local activity at the spatial scale of the distance between the anode and cathode (approximately 1 cm), but can also alter more distant regions. Because memory relies on a broad network of cortical and subcortical regions, including the hippocampus (Kim, 2011; Keerativittayayut et al., 2018), stimulating a broader network may be necessary to impact cognitive function. On the other hand, memory also relies on the recapitulation of specific patterns of neuronal activity, especially within the hippocampus (Foster, 2017; Staresina and Wimber, 2019). Thus, other work has stimulated through microelectrodes to mimic and reinstate memory-related hippocampal activity using a model-based closed loop approach (Hampson et al., 2018; Hampson et al., 2013; Deadwyler et al., 2017). An avenue for future work could use macroelectrode stimulation in a similar vein, by triggering stimulation at multiple macroelectrode contacts in order to synchronize a particular spatiotemporal pattern of activity across key memory-related regions (Kim et al., 2016; Kim et al., 2018).

In relating low-frequency network connectivity, physiology, and behavior, our study contributes to methodological development for invasive stimulation (Krauss et al., 2021; Cagnan et al., 2019) that illuminates the critical role of low-frequency networks in cognition (Voytek and Knight, 2015). In addition, the present study also suggests that other methods that manipulate low-frequency activity could be leveraged to modulate neural and cognitive function. Several recent studies using non-invasive methods have leveraged low-frequency theta-patterned stimulation to modulate episodic and working memory (Nilakantan et al., 2017; Hermiller et al., 2020; Tambini et al., 2018; Warren et al., 2019; Grover et al., 2022). Such low-frequency stimulation modulates electrophysiology perhaps by entraining low-frequency oscillations that are associated with cognitive function (Solomon et al., 2021; Reinhart and Nguyen, 2019; Reinhart et al., 2017; Hanslmayr et al., 2019).

In summary, our demonstration of improved memory with closed-loop stimulation supports the idea that memory function is dynamic, and that closed-loop algorithms that account for moment-to-moment variability in the brain’s memory state can selectively deliver stimulation only when it is needed. The present study also links closed-loop stimulation efficacy to white matter targeting, brain-wide evoked physiology, and changes in episodic memory performance. The findings suggest future strategies for using the functional and anatomical network profile of putative stimulation targets to optimize downstream changes in oscillatory activity and cognition.

## Acknowledgments

This work was supported by NIH grant NS106611 and MTEC project 20-06-MOM from the Army Medical Research and Development Command. We are indebted to the patients and their families for their participation and support. The views, opinions, and/or findings contained in this material are those of the authors and should not be interpreted as representing the official views or policies of the Department of Defense or the U.S. Government. B.C.J. receives research funding from NeuroPace and Medtronic not relating to this research. M.J.K. and D.S.R. each hold a greater than 5% equity interest in Nia Therapeutics, LLC, a company intended to develop and commercialize brain stimulation therapies for memory restoration.

## Notes

### Competing Interest Statement

The authors have declared no competing interest.

### Summary of Updates

Revisions to various sections of the text, including the Abstract, Introduction, and Methods.

